# Urbanisation and habitat loss favour thermophilic and monogynous ant species

**DOI:** 10.1101/2025.02.25.640022

**Authors:** Basile Finand, Thibaud Monnin, Céline Bocquet, Angélique Bultelle, Pierre Fédérici, Léa Darmedru, Solène Gouffault, Joséphine Ledamoisel, Nicolas Loeuille

**Affiliations:** Sorbonne Université, Université Paris Cité, Université Paris Est Créteil, CNRS, INRAE, IRD, Institut d’Ecologie et des Sciences de l’Environnement de Paris (UMR7618), 75005 Paris, France; Ecosystems and Environment Research Program, Faculty of Biological and Environmental Sciences, University of Helsinki, Helsinki, Finland

**Keywords:** Ants, heat island effect, metacommunity, trait selection, urbanisation

## Abstract

Environmental changes such as urbanisation and related habitat loss and fragmentation profoundly impact ecological communities by altering habitats, resources, and microclimates. Yet, the impacts of the resulting environmental conditions on metacommunity dynamics, ranging from species sorting to mass effects, remain a subject of debate. Ants, with diverse life histories and strong ecological effects, are ideal model species to study these pressures. We investigated the response of ant communities, including taxonomic and functional diversity, to urbanisation and habitat fragmentation in the Paris region, comparing wooded areas in 25 urban parks vs. 24 rural forests outside the city. We found a clear difference in species composition between urban and rural environments, with a higher prevalence of thermophilic species and a tendency for monogyny in the city. Forest communities were homogeneous across the size levels we studied, while park communities differed noticeably depending on park size, with larger parks harbouring more species. Our findings suggest that urbanisation selects specific ant traits and favours more thermophilic species, thereby increasing the mean thermal preference of urban communities. These selective effects influence which species can colonise and survive in different patches, shaping metacommunity structure and potentially affecting the resilience of ant communities under climate change.

## Introduction

Habitat loss, and the fragmentation it often entails, stand as a primary driver of biodiversity decline, exerting manifold impacts on ecosystems worldwide (Alberti, 2015; Aronson et al., 2014; Diamond & Martin, 2021; Haddad et al., 2015; Johnson & Munshi-South, 2017; Piano et al., 2020). Habitat fragmentation is the process by which a once-contiguous habitat is progressively broken into a greater number of patches, smaller and more isolated from one another than the initial habitat, together with an overall reduction in habitat amount (Fahrig, 2003). The resulting fragmented habitats are therefore characterised, at a given point in time, by a landscape of numerous, small, and isolated patches within a reduced total habitat amount. An ongoing debate separates the effects of habitat fragmentation “per se” (ie, for a fixed amount of habitat) versus habitat loss (Fahrig, 2017; Fahrig et al., 2019; Riva et al., 2024). Despite the interest in understanding the effects of these two factors, when considered over time, habitat fragmentation is often a consequence of habitat loss, and these processes are therefore highly related. Therefore, hereafter, we use habitat fragmentation to refer to fragmented habitat distributions, resulting from both habitat loss and fragmentation, and we characterise it based on patch size distribution, relative isolation and connectivity.

Habitat fragmentation impedes dispersal between populations, leading to direct impacts on individuals and gene flows, resulting in population declines and reductions in genetic variability (Hastings, 1983; Travis et al., 2013; Young et al., 1996). These effects reverberate through ecological communities, altering their composition and dynamics. Habitat fragmentation gives rise to metacommunities, i.e. interconnected communities linked through dispersal. Metacommunity dynamics can happen in various ways depending on species responses to environmental characteristics, biotic interactions, dispersal, and the stochasticity of these processes. This leads to four paradigms (Leibold et al., 2004). First, in scenarios where variations in environmental conditions among patches do not play a key role, biodiversity hinges exclusively on the competitive abilities and dispersal capacities of species, a dynamic termed patch dynamics. Dispersing species can exploit and establish populations in patches where competitors cannot disperse (Tilman, 1994). In this “patch dynamics” paradigm, habitat distributions made of small and isolated patches are expected to favour dispersing species while disadvantaging more competitive species, thereby reducing total diversity (Finand, Monnin, et al., 2024; Tilman et al., 1994). Second, in environments where variability among patches drives ecological dynamics and where dispersal is restricted, species diversity is governed by species sorting (second paradigm), where species whose ecological niches best match local conditions competitively exclude others within each patch. Third, in environments where variability among patches drives ecological dynamics and where dispersal between patches is extensive, mass effects (third paradigm) come into play (Mouquet & Loreau, 2003), where spatial fluxes of individuals dominate the local species sorting process. Less-adapted species persist in small numbers because they are continually resupplied from surrounding patches where they are better adapted, due to source-sink fluxes. However, when dispersal is exceedingly high and mass effects dominate, all species compete for average metacommunity conditions. This leads to competitive exclusion at this scale, and diversity becomes low at both local and metacommunity scales (Mouquet & Loreau, 2003). Heightened fragmentation of habitat, by augmenting patch isolation, is expected to foster species sorting, consequently diminishing local species richness and increasing species turnover among patches. Last, the fourth paradigm, the neutral theory, akin to island biogeography (MacArthur & Wilson, 1967), posits that diversity is solely dependent on species loss through emigration and extinction balanced by gains through immigration and speciation, with no differentiation in species niche, competitive interactions, and dispersal abilities playing a role in the process (Hubbell, 2001).

Importantly, these four paradigms can be disentangled based on variations in species composition and patterns of local (alpha), regional (gamma) diversity and variations in species composition among patches (beta diversity) (see e.g. Leibold & Mikkelson, 2002). For instance, if species sorting dominates, local composition should differ in contrasted environments so that beta diversity is mostly associated with a “turnover” component (sensu Baselga, 2010). Conversely, when dispersal leads to mass effects, we expect a spatial autocorrelation in species composition so that communities in harsher environments or smaller patches should be a subset of communities developing in better or larger patches. This enhances the “nestedness” component of beta diversity (sensu Baselga, 2010). If considering the “patch dynamics” paradigm, species in more fragmented environments (e.g., within cities) should be a subset of other communities (leading to nestedness), this subset being explained by competition-dispersal abilities, with more dispersive species favoured in fragmented contexts (Tilman et al., 1994).

Urbanisation is another major factor affecting biodiversity. This transformative process reshapes environments in many ways, fostering novel ecological dynamics and posing significant challenges to native flora and fauna. First, the urbanisation process involves habitat destruction, which results in low habitat amount and fragmented distributions of habitat patches within the urban matrix. Second, it also generates varied types of pollution, including chemicals, noise disturbance, and altered light regimes, often favouring species with specific adaptations or tolerances (Hölker et al., 2010; Newport et al., 2014). These pollutants permeate air, water, and soil, exerting deleterious effects on biotic components of ecosystems. Third, changes in resource availability represent another hallmark of urbanisation, including variations in quantity, quality, and seasonality of food resources (Hantak et al., 2021; Wilson & Jamieson, 2019). Finally, the creation of distinct microclimates within cities, characterised by elevated temperatures (urban heat island effect), exacerbates thermal stress on organisms and modifies ecosystem dynamics (Kaiser et al., 2016; McGlynn et al., 2019). This thermal gradient can influence species distributions, phenology, and physiological processes, driving community composition and structure shifts.

Beyond diversity indices, the specific environmental factors and the low connectivity encountered in cities should select specific traits potentially leading to trait syndromes (Hahs et al., 2023). Habitat fragmentation may favour low dispersal (Cheptou et al., 2008; Finand et al., 2023). In a spatially heterogeneous but temporally stable environment, selection disfavours dispersal, since it relocates individuals away from patches they are already optimally suited to, with no temporal variation to justify the move (Hastings, 1983). Conversely, better dispersers can be selected in fragmented contexts due to competition-colonisation trade-offs, where dispersal allows access to empty patches ahead of superior competitors (Finand, Monnin, et al., 2024; Tilman et al., 1994). Higher temperatures promote species and phenotypes whose thermal preference matches these conditions (Campbell-Staton et al., 2021; Diamond et al., 2018). For instance, longer-legged ant species are selected in hotter parks in Taichung City because the distance between the body and soil limits desiccation (Liu et al., 2019). In general, urban and rural species may differ in many other traits (Buchholz & Egerer, 2020; Croci et al., 2008; Hahs et al., 2023), reflecting the multidimensional changes in such environments.

Insects, and in particular ants, are a very good model group to study the effects of urbanisation and habitat fragmentation on communities. They have important impacts on ecosystems, dispersing seeds, pollinating, participating in nutrient cycles, structuring the soil and contributing to food chains (Hölldobler & Wilson, 1990). First, their life history traits are variable between species. Some of these traits are strongly related to their dispersal abilities (size of individuals, presence or absence of wings in queens) and competitiveness (size of colonies, diet), and therefore play a key role as functional traits in metacommunity dynamics. Second, ants are abundantly present and diverse inside and outside cities, which allows direct comparative studies. Most ant species depend on green spaces, which makes the city a very fragmented habitat compared to the countryside, even if some species can live in built areas. Last, they are easy to sample. Urbanisation and habitat fragmentation have been shown to decrease species richness in ants and to change the composition of their communities (Carpintero & Reyes-López, 2014; Ješovnik & Bujan, 2021; Leal et al., 2012; Pacheco & Vasconcelos, 2007; Theunis et al., 2005). For instance, climate and resource specialist ants are more sensitive to habitat fragmentation (Leal et al., 2012). Moreover, this change in community composition due to habitat fragmentation impacts ecosystem services provided by ants, influencing decomposition and soil structure (Sanford et al., 2009).

We here investigate the impact of urbanisation and habitat fragmentation, together with local-scale variation in habitat amount and isolation, on ant communities at different scales by comparing forest (low urbanisation and low fragmentation) and urban communities in wooded areas of parks (high urbanisation and high fragmentation). Paris and its region offer a good opportunity to study urbanisation and habitat fragmentation, with a marked gradient from the very densely populated city centre to low-density areas thirty kilometres away. The city has numerous urban parks with wooded areas of various sizes, surrounded by a matrix of streets and buildings unsuitable for most ant species. This makes Parisian parks a perfect setup to understand the impact of fragmentation on ant communities. Moreover, the urban heat island effect within Paris is well-documented, on average 2–3°C warmer than surrounding rural areas annually, with nocturnal differences reaching up to 6°C (Cantat, 2004; Lemonsu et al., 2015). The surrounding region is more rural, with many natural forests of various sizes, allowing an interesting comparison with the city. We measured community differences between the two habitats, and the effects of local-scale habitat distribution, isolation and habitat amount within urban parks, and habitat amount within forests. We hypothesise that diversity is lower in parks due to their smaller size and higher isolation from other patches in fragmented urban habitat settings. Indeed, ant dispersal is limited, especially in urban environments. Within parks and within forests, we expect a decrease in species richness in smaller and more isolated habitats due to lower mass effect. Within parks, this decrease should be more intense due to additional dispersal barriers. Finally, we studied variations in traits linked to thermal preference, colony social structure (queen number) and dispersal abilities. We hypothesise that the city harbours species that are more heat-resistant due to the urban heat island effect, and that have short dispersal distances because of the high habitat fragmentation.

## Material and methods

### Patch selection

Litter ants depend on wooded habitats, where leaf litter and canopy cover provide the moisture, nesting sites, and food resources they require. To study the impact of urbanisation and habitat fragmentation on litter ants, we sampled two highly contrasted habitats: 24 rural forests in the Paris region and wooded parts of 25 urban parks within the city (Figure 1). We stress that this design does not disentangle the effect of habitat fragmentation from other environmental changes associated with urbanisation, such as temperature, pollution, or resource availability. Forests were located in minimally fragmented environments, as they offer a favourable habitat for litter ants and are surrounded by a matrix of habitats that, while less favourable, can still support ant species, e.g. hedges around agricultural fields or private gardens. On the contrary, parks are islands of favourable habitats (woods and shrubs) surrounded by a mostly hostile matrix (streets and buildings), though local refugia may exist. To compare the same litter ant communities in forests and parks, we targeted park areas with natural litter for sampling, i.e., areas under trees or shrubs. This allowed us to focus on the specific community of ants living in the litter and to make habitats as comparable as possible. Forests adjacent to the city were used here as reference communities to assess the effect of urbanisation, which intrinsically involves habitat change. Differences between park and forest habitats reflected spatial variations in niche conditions, which is precisely what this study aimed to examine. We studied the effect of urbanisation at the regional scale (city parks vs. forests) and the effect of different components of habitat fragmentation at the local scale (parks of varied size and isolation, forests of varied size). At local scales, we stressed that habitat amount is the key factor. While heterogeneity within patches definitely plays a role in the maintenance of diversity, its quantification would ultimately need information on ecological niches that is not available for most species. Our study did not aim to study the role of this intra-patch heterogeneity.

**Figure 1:**
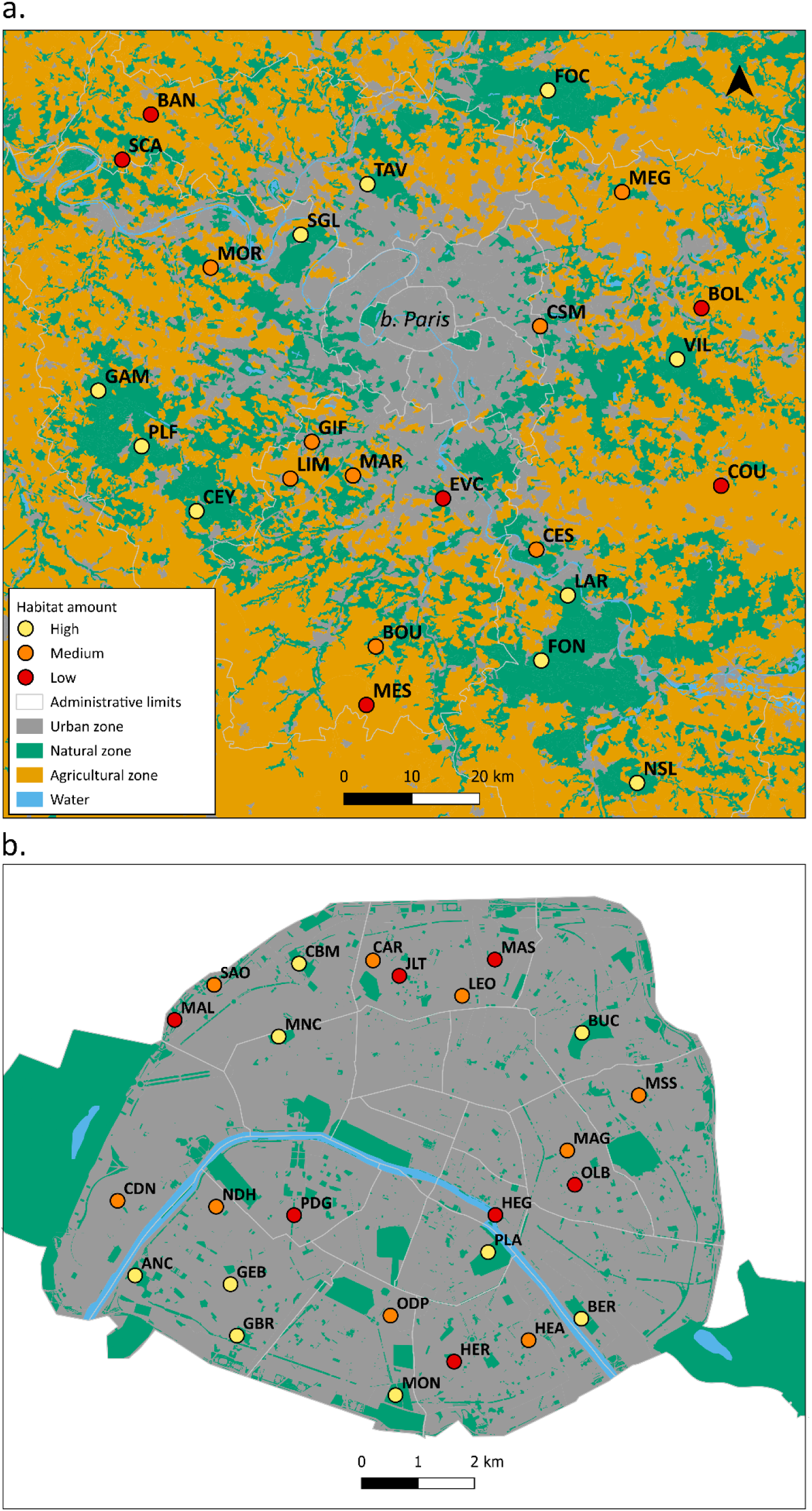
Map of the sampling sites in forests around Paris (Ile-de-France region) (a) and parks in Paris (b). The colour of the circles indicates the habitat amount. High habitat amount forests (n = 10) have between 75 and 95% of forest cover within 1 Km, medium habitat amount forests (n = 8) have between 30 and 70% of forest cover, and low habitat amount forests (n = 6) have less than 5% of forest cover in their surroundings. Parks are divided based on surface: high habitat amount (80,000 to 250,000 m² (n = 8)), medium habitat amount (5,000 to 12,000 m² (n = 9)), and low habitat amount (500-2,000 m² (n = 8)). Site information is detailed in Table S1. Background data are extracted from the Corine Land Cover.

We selected deciduous forest patches (n = 24) of three habitat amount classes (Figure 1a). High habitat amount forests (n = 10) had between 75 and 95% of connected forest cover within a 1 Km radius centred on the sampling point, medium habitat amount forests (n = 8) had between 30 and 70% of connected forest cover, and low habitat amount forests (n = 6) had less than 5% of connected forest cover in their surroundings. We used the amount of habitat within a 1 Km radius, which is coherent with the mean dispersal distance of ants (Helms, 2018; Kuhn et al., 2017; Suni & Gordon, 2010). In addition, using the total size of the forests would not make sense as most were too large (tens to hundreds of Km²) compared to the ant’s scale. We used the database BD Forêt® Version 2.0 from the “National Institute of Geographic and Forest Information” of France (IGN, https://inventaire-forestier.ign.fr/carto/afficherCarto/V2, resolution: 50 cm, last visit: 19/07/2022) to assess forest cover. Forest classes were evenly distributed in the Paris region, and we chose well-separated locations that likely represent independent plots (mean distance = 52.1 ± 23.7 Km, min = 6.4 Km, max = 122.1 Km).

Local-scale habitat distribution of parks was characterised using two metrics, size (habitat amount) and isolation. We selected parks (n = 25) of three size classes representative of the whole distribution of park size in Paris with a homogeneous representation of the different classes (Figure 1b), namely high habitat amount (n = 8, 80,000-250,000 m²), medium habitat amount (n = 9, 5,000-12,000 m²), and low habitat amount (n = 8, 500-2,000 m²). We calculated the proximity index as a measure of isolation (Gustafson & Parker, 1994). It accounted for the quantity of green area (other parks) within 1 Km around the focal park boundaries (expected mean dispersal distance of ants; Helms, 2018; Kuhn et al., 2017; Suni & Gordon, 2010), with weights directly related to the distance between patches. As for forests, we selected sites that were evenly distributed within Paris and well separated, assumed to represent independent points (mean distance = 4.9 ± 2.1 Km, min = 552 m, max = 9.5 Km). All parks were managed by the City of Paris, indicating similar management, except PLA, which is managed by the French National Museum of Natural History. We used the City of Paris dataset of green spaces (https://opendata.paris.fr/explore/embed/dataset/espaces_verts/map/), and we added manually the green spaces not managed by the City of Paris, such as the “Jardin des Plantes”, the “Cité Internationale Universitaire de Paris”, or the “Jardin des Tuileries”.

### Sampling

We sampled forests between May and July 2020, between 11 am and 4 pm, on rainless days. We distributed 12 quadrats of one square meter each, equally spaced every 9 m, along a transect of 100 m. We independently collected the litter and superficial soil on each of the 12 quadrats. We collected organisms using the Winkler extraction method (Agosti et al., 2000). We detail the procedure below.

We sampled parks between May and August 2021, between 11 am and 4 pm, on rainless days. To compare the same litter ant communities in forests and parks, we targeted park areas with natural litter for sampling, i.e. areas under trees and shrubs (always when litter was present). It allowed us to focus on the specific community of ants living in the litter and make the habitats as comparable as possible with forests. We purposefully placed the 12 quadrats into these areas and dispersed the quadrats within parks as much as possible (considering the configuration of the parks, transect was not possible. Minimum distance between the quadrats: 9 m). One park (OLB) was so small that only five quadrats could be placed and sampled, and owing to a disturbance only ten quadrats were sampled in another park (MAS). The sampling and extraction protocols were the same for parks and forests.

For each quadrat, we sifted the litter and the first centimetres of soil using a 1 cm² sieve (Figure S1a). This removed leaves, twigs and stones and retained the soil and organisms within, including ants. We kept what passed the sieve in large plastic bags until we returned to the laboratory. There, we moved the litter to sieve bags that were hung in a Winkler extractor for 48 hours (Agosti et al., 2000) (Figure S1b). As the litter gradually dried out, organisms moved away and fell into an alcohol-filled collecting vial. This method samples all the macrofauna living in the collected soil and litter, including ant foragers present in the litter and parts of the colonies that nest in the litter or superficially in the soil.

We sampled litter ants in forests and parks in two separate years (2020 and 2021) due to logistical constraints and movement restrictions related to the COVID-19 pandemic. Note that ant queens and colonies live several years, and that the sampling was done during the same season. While fluctuations of abundances from one year to another are certainly possible, our analyses focus on species presence/absence in each quadrat rather than abundance. We further discuss the implications of our sampling scheme in the discussion section.

### Identification of ant species

We identified ants at the species level using a binocular microscope and three identification keys (Blatrix et al., 2013; Seifert, 2018; Galkowski & Lebas “Identification des *Myrmica*”, unpublished). In addition, *Temnothorax, Hypoponera* and *Tetramorium* species were identified with the help of experts (Xavier Espadaler for the two first groups, Bernard Kaufmann for the last).

### Ant species traits

We selected different species-level traits to test for the effect of urbanisation. Queen size, worker size and colony size (number of individuals) were obtained from the bibliography for all collected species (Blatrix et al., 2013; Seifert, 2018). In addition, colony mass was estimated as worker size^3^ × colony size. Because colony mass is directly linked to metabolic requirements (Brown et al., 2004), we expected that it could vary between urban and forest sites, either due to changes in resources or temperature regimes.

We also obtained the social structure of a colony (Blatrix et al., 2013; Seifert, 2018), with two levels: monogyny (species with typically one queen per colony) and polygyny (species with typically several queens per colony). This trait is to some extent linked to dispersal as monogynous species often perform independent colony foundation (long dispersal distance strategy but low competitive ability), whereas polygynous species do colony fission (short dispersal distance strategy but high competitive ability) (Cronin et al., 2013; Keller, 1991).

Finally, for each species, we estimated thermophily based on a meridional index from Daufresne et al. (2004). Recent advances in ant thermal ecology have highlighted more precise traits such as critical thermal limits, activity temperatures, or behavioural tolerances (Nascimento et al., 2022; Parr & Bishop, 2022; Roeder et al., 2021). They are well-suited to characterising precisely the thermal performances of a few species in a given environment. However, they require live workers in significant numbers (20 to 30 workers per colony, more for polymorphic species, and several colonies per species), making them incompatible with our sampling protocol with Winklers. The meridional index is better suited for analysing diverse assemblages across broad spatial scales, like in our study. In addition, it integrates factors other than thermal resistance per se that may relate to thermal preference, e.g. circadian rhythm of activity, and it is generalisable across taxa and compatible with studies on biogeographical shifts under climate change. It provides a useful, scalable metric for detecting broad patterns of community shifts along environmental gradients. Species were considered meridional if most of their distribution lies south of Paris, with the meridional index being calculated by (Paris latitude – minimum observed latitude) / (maximum observed latitude – minimum observed latitude). The meridional index for a given species is, therefore, close to one when Paris is at the northern limit of its distribution, and close to zero when Paris is at the southern limit of its distribution. Minimum and maximum latitudes were obtained for each species distribution from the antmap.org website (last visit: 28/02/2024; Guénard et al., 2017; Janicki et al., 2016). Only the records considered to pertain to the native distribution were considered. Following previous works, we used this index as a proxy of the thermal preference of a species. A Parisian species at the northern limit of its distribution is more thermophilic than one at the southern limit. Paris is farther north than the northern range limit for two species. We have attributed 1, the highest value, to them. Removing these species did not change the results.

To understand which traits are linked to urbanisation, we assigned a value of urbanisation preference to each species, calculated as the number of urban patches where the species has been found divided by the total number of patches (urban and forest patches) where it has been found. Strictly urban and forest species have urbanisation indices of one and zero, respectively.

### Analyses

Statistical analyses were done with R v4.5.2 (R Core Team, 2022). We checked that our sampling was sufficient to sample the litter ant biodiversity with rarefaction curves and Chao analysis using the vegan package.

First, at the regional scale, we analysed alpha diversity (species richness and Shannon index) as a function of urbanisation (forests vs. parks) using linear models. We focused on species richness because it was highly correlated with the Shannon index (R² = 0.92; but see Figure S5 for the Shannon diversity analyses). Second, at the local scale, we focused on community variations among patches of the same habitat type. In parks, we analysed species richness and the Shannon index as a function of park size (habitat amount) and isolation using an LM. In forests, we analysed species richness and the Shannon index as a function of habitat amount, using a generalised linear model with a Poisson distribution, as residuals were not normally distributed. For all models, we checked that the data complied with application conditions using the DHARMa package (Hartig, 2022).

We assessed beta diversity with Jaccard’s dissimilarity index because we focused here on species presence/absence rather than on abundances. Using individual densities would introduce biases because ants are social species, and colony sizes differ among species. Our sampling procedure can recover isolated individuals (i.e. independent samples) as well as groups of foragers or even whole colonies for some *Temnothorax* species and *Myrmecina graminicola* (non-independent samples). Large groups or colonies would inflate recorded abundances of superficial species compared to species nesting deeper in the soil, for which we only collected foragers. We performed db-RDA (distance-based redundancy analysis), an RDA on dissimilarity distances, to assess whether our environmental factors affect dissimilarity between our communities (Legendre & Anderson, 1999). We used linear models to test the significance using the vegan package, and the package *betapart* to assess the relative contribution of turnover and nestedness composing the dissimilarities between our communities (Baselga & Orme, 2012).

To link urbanisation preference to species traits, we first tested the correlation among traits using a Pearson test. The different size measurements were highly correlated (Figure S2). We kept the total mass of a colony, as it contains the size of the colony and the worker size. The influence of social structure, colony mass, and meridional index on urbanisation preference was analysed using Generalised Linear Models (GLMs, Douma & Weedon, 2019). Given the nature of the response variable (proportions derived from site counts), we employed a Beta-binomial distribution to account for overdispersion using the glmmTMB package (Brooks et al., 2017). A model selection procedure based on the Akaike Information Criterion (AIC) was performed to compare the fit of various candidate models, ranging from a null model to a full global model. To further explore the individual contribution of each trait and to account for potential limitations in statistical power due to our sample size, we analysed univariate beta-binomial regressions for each predictor separately and the full model. Multicollinearity among predictors was assessed using the Variance Inflation Factor (VIF) via the performance package (Lüdecke et al., 2021), ensuring that all included variables met the requirement of VIF < 2. Model validation was conducted by inspecting simulated residuals using the DHARMa package (Hartig, 2022). Colony mass was log-transformed. All figures represent marginal effects calculated with the ggeffects package (Lüdecke, 2018).

## Results

We collected and identified 36,361 ants from 12 genera and 29 species. Abundances were similar in forests and parks (18,094 and 18,267 ants, respectively). Similarly, the number of species recovered per habitat type (gamma diversity) was comparable (Figure 2, 23 species in forests and 20 species in parks). 14 species occurred in both parks and forests, albeit in most cases with different relative frequencies, while 9 species occurred in forests and 6 in parks only (Figure 2). Only one exotic species was recorded (*Lasius neglectus*). Removing it from the analyses did not qualitatively change the results. The high prevalence of *Temnothorax nylanderi* and *Myrmecina graminicola* reflects that our samples were representative of the forest litter ant communities. These two species, which dominated our park samples, are characteristic forest dwellers (Blatrix et al., 2013; Seifert, 2018) and were also the most abundant ants in our forest sites.

**Figure 2:**
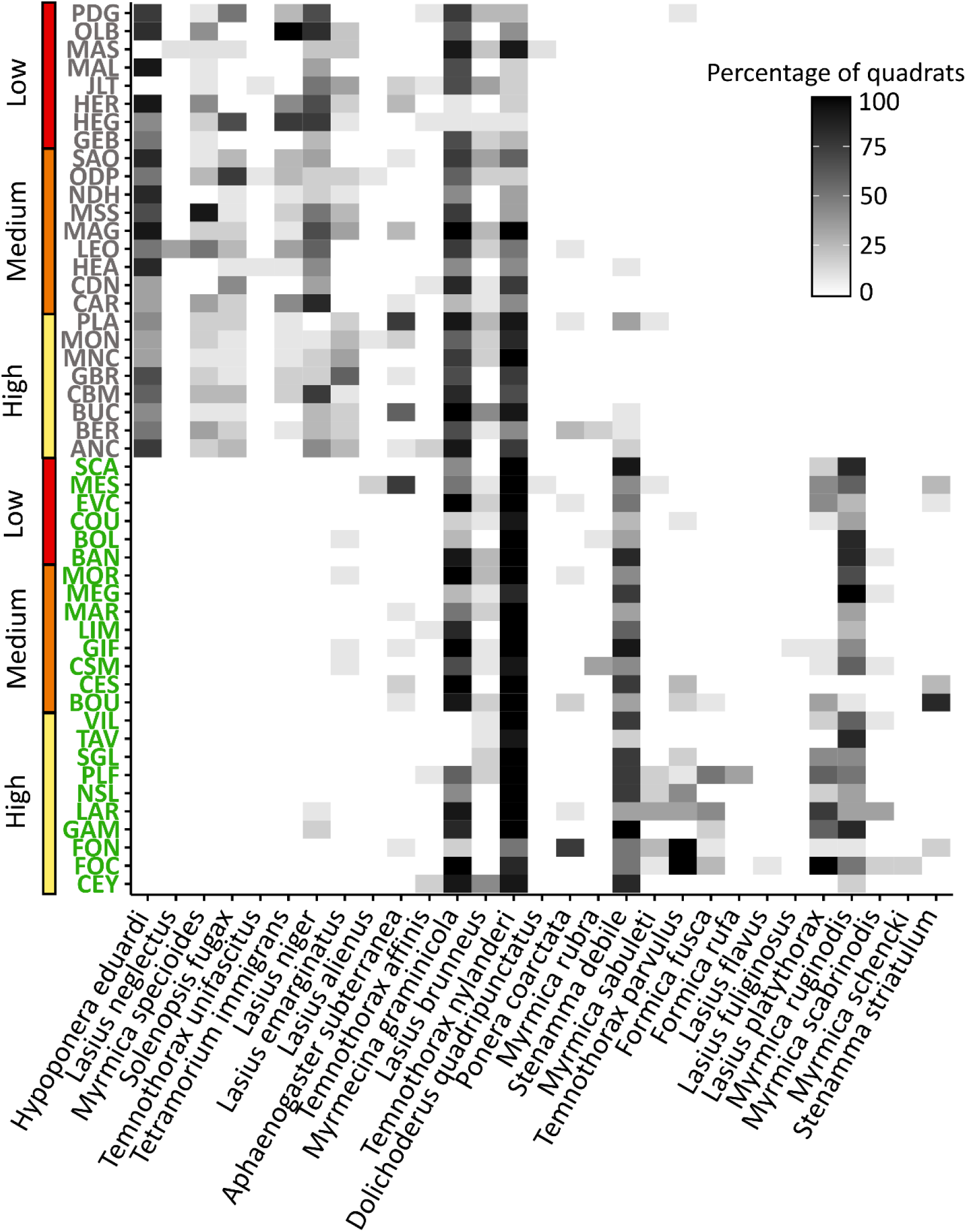
Variations in the presence and frequency of ant species depending on patch types. Darker Levels show higher percentage of quadrats where the species was found. On the Y-axis, green patches correspond to forests and grey patches correspond to parks. Red, orange, and yellow show habitat amount levels. Species are ranked based on their urbanisation preference along the X-axis and alphabetically for those with the same value (i.e. absent in one habitat).

### Sampling effort

The rarefaction curves and the Chao analyses showed that the sampling was sufficient to estimate litter ant diversity (Figure S3). Indeed, we sampled 95.19 ± 25.41% of the theoretical number of species in forests and 91.13 ± 13.12% in parks. We sampled less than 80% in only two forests (CSM and GIF), and three parks (HEA, HEG and PDG). However, this theoretical number of species was higher in these patches (around or above 15) than in the other patches (usually around 10, seldom higher than 12), so it may have been overestimated. Therefore, we used the observed number of species per patch in the analyses.

### Effect of urbanisation at the regional scale (forests vs. parks)

Unexpectedly, we found more species in urban parks (8.96 ± 1.74 species) than in forests (7.58 ± 2.30 species, F value = 5.6, p = 0.022, R² = 8.7%, Figure 3a).

**Figure 3:**
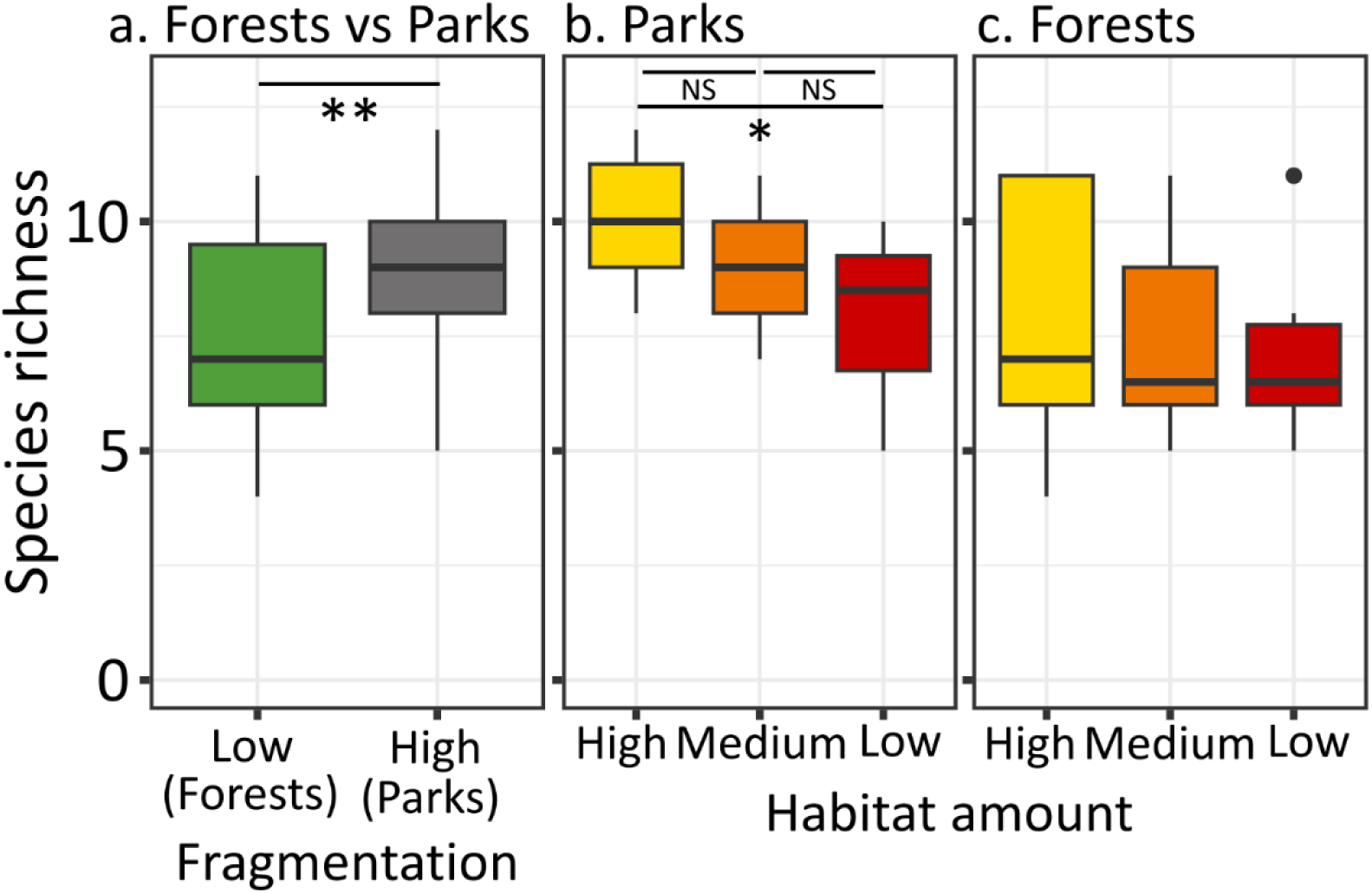
Species richness between forests and parks (a), between parks of different habitat amount (b), and between forests of different habitat amount (c). Significant differences are observed in species richness between forests and parks (F value = 5.6, p = 0.022, R² = 8.7%), and in species richness between parks of different habitat amounts (F = 3.74, p = 0.039, R² = 0.19).

Urbanisation also influenced species composition (F value = 28.47, p = 0.001, R² = 37.7%, Figure 4a). Communities were clearly different between parks and forests, with 89% of the beta diversity due to turnover (Jaccard index = 0.517, Figure 2), suggesting a dominant role of species sorting.

**Figure 4:**
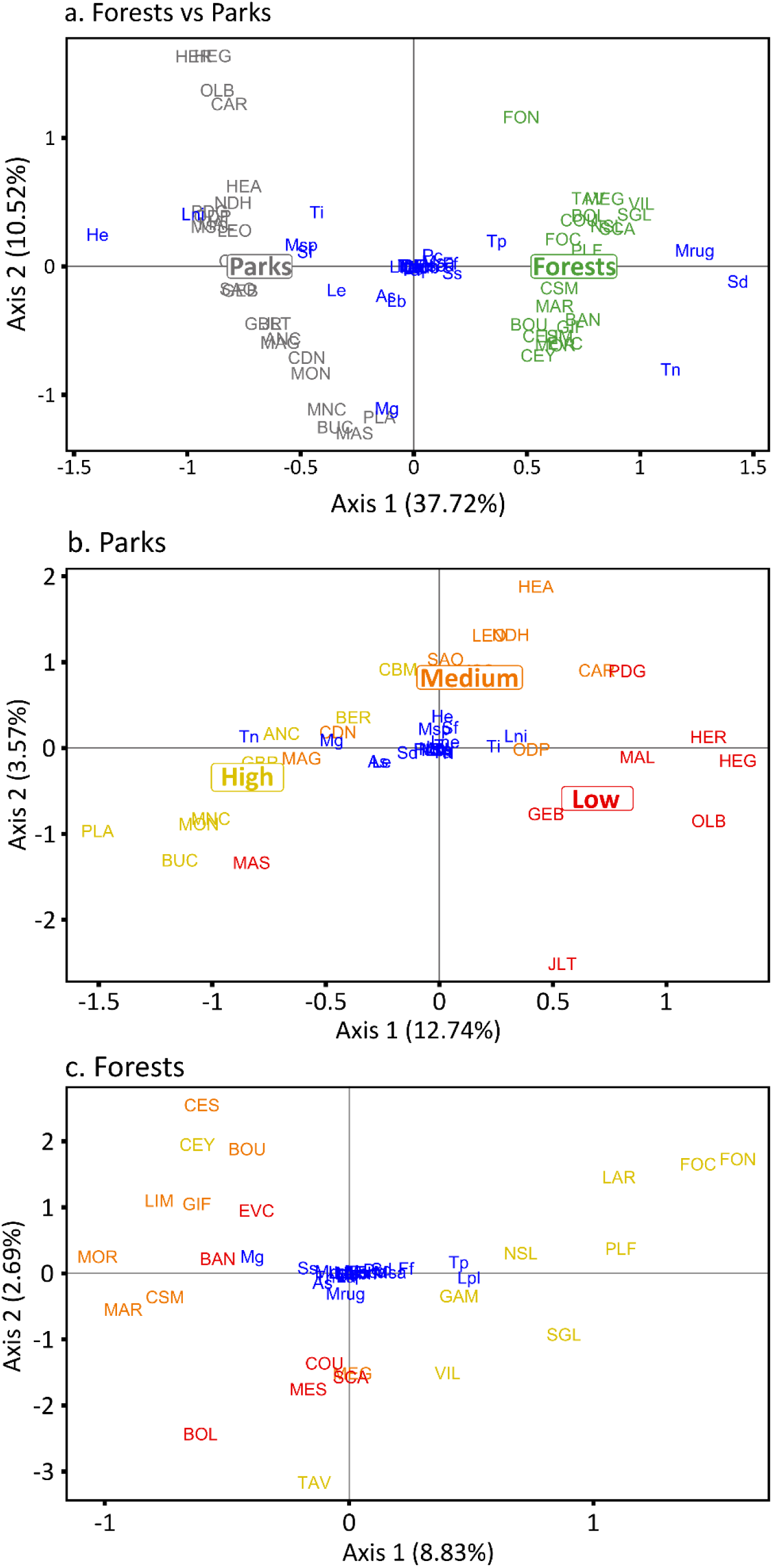
Beta diversity (db-RDA analysis) between forests and parks (a), between parks (b), and between forests (c). Blue acronyms are the species. Capital acronyms are the sites. Yellow represents high habitat amount, orange medium habitat amount, and red low habitat amount. Ant communities are markedly different between parks and forests, with communities from the two habitats being separated with no overlap (F value = 28.47, p = 0.001, R² = 37.7%). Communities also clearly differ between parks of different habitat amount, with a gradual change in composition from low to medium to high habitat amount (F value = 2.14, p = 0.006, R² = 16.3%). No difference is observed between forests of different habitat amount. (F = 1.37, p = 0.12).

### Effect of habitat amount and isolation at the local scale

In the forests, habitat amount had no effect on species richness (Chisq = 0.411, p = 0.81, Figure 3c) nor on the composition of ant communities (F value = 1.37, p = 0.12, Figure 4c).

In contrast to forests, the habitat amount had an effect on species richness and community composition in parks. As expected, in the city, larger parks harboured more species than smaller parks (F = 3.74, p = 0.039, R² = 0.19, Figure 3b). The effect was significant between small and large parks (pairwise comparison: large-medium p = 0.21, medium-small p = 0.57, large-small p = 0.03). High habitat amount (large) parks had on average 10.1 ± 1.5 species, medium parks had 8.8 ± 1.4 species and small ones had 8.0 ± 1.9 species. Habitat amount also affected the spatial turnover of ant communities (F value = 2.14, p = 0.006, R² = 16.3%, Figure 4b). The beta diversity was composed of 100% of turnover between large and medium parks, 84% between medium and small, and 93% between large and small (Jaccard index large-medium = 0.222; medium-small = 0.278; large-small = 0.450), highlighting a dominant role of species sorting.

For parks, isolation did not affect species richness (t value = -1.63, p = 0.12, Figure S4a), or ant communities composition (F = 0.76, p = 0.70, result not shown).

### Correlation between species traits and urbanisation preference

According to the multivariate model, urbanisation preference was not significantly influenced by colony mass (Chisq = 0.09, p = 0.76, Figure 5a) or social structure (Chisq = 0.99, p = 0.32, Figure 5b). However, urbanisation preference was significantly correlated with a higher meridional index (Chisq = 4.37, p = 0.037, Figure 5c), suggesting that urban environments select for more thermophilic ant species. While univariate models yielded similar results for most traits, a marginal trend emerged for social structure, the urban context tending to favour monogynous species (Chisq = 3.48, p = 0.062).

**Figure 5:**
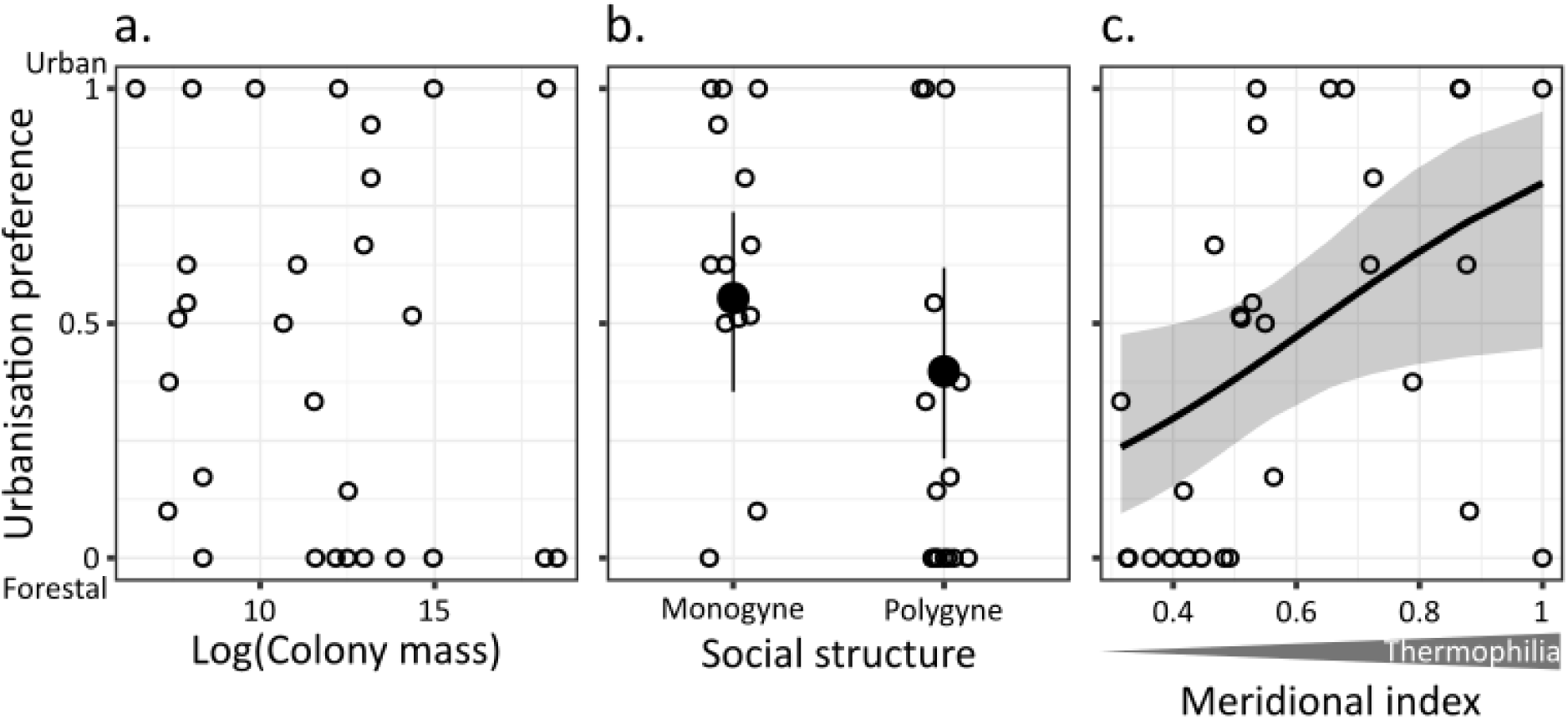
Relation between Colony mass (a), Social structure (b), Meridional index as a proxy of thermophily (c) and the urbanisation preference. Urbanisation preference varies between zero for strictly forest species and one for strictly urban species. The colony mass has no significant effect (Chisq = 0.09, p = 0.76). The social structure has a marginal effect on urbanisation preference in the univariate model (Chisq = 3.48, p = 0.062). The Meridional index is correlated to urbanisation preference (Chisq = 4.37, p = 0.037). Open circles represent the raw observed proportions for each species. For (b), the black dots represent the predicted marginal means for the proportion of urban sites for each social category (Monogyne vs. Polygyne), with error bars indicating the 95% confidence intervals. Open circles have a horizontal jitter for clarity, maintaining their exact values on the y-axis. For (c), the solid black line represents the predicted proportion of urban sites as a function of the latitude index, derived from the Beta-binomial GLMM. The shaded area indicates the 95% confidence interval.

## Discussion

Our study showed that urbanisation and the associated habitat fragmentation influenced the litter ant communities at the metacommunity scale. The ant biodiversity assessed by the total number of species per environment (gamma diversity) was stable, and the species richness per patch was slightly higher in parks than in forests. Yet, communities differed markedly between the two environments, with a strong turnover in species compositions. At a local scale (alpha diversity), species richness and community composition were affected by habitat amount in parks and did not vary among forests. Finally, we highlighted that the turnover in species composition between urban and forest patches was associated with systematic differences in species phenotypes. Urbanisation selected species that were thermophilic, and there was suggestive evidence that it also favoured monogynous species.

### Methodological considerations

A possible limitation of our study is that forests and urban parks were sampled in different years (2020 vs. 2021). While we acknowledge this issue, we however think that potential biases are limited here. First, our method relies on community composition (presence/absence of species), not on species abundances. While abundances likely fluctuate from a year to the next, such variations are not expected when looking at the presence/absence of long-lived species. Ant queens and colonies are indeed long-lived (Hölldobler & Wilson, 1990). Ant queens live on average 10 years, up to 28 years, and colonies outlive queens in species where colonies have several queens or where queens can be replaced, e.g. by adoption of daughter queens (Keller, 1998). The composition of ant species community is therefore relatively stable from one year to another (Donoso, 2017; Herbers, 2011). While we cannot exclude that individual species go extinct or colonise a patch between the two years, it is unlikely to affect the broader scale diversity pattern and strong turnover components that we observe here. Focusing on species presence/absence therefore reduces sensitivity to yearly fluctuations in colony size or activity. The clear differences observed between forests and parks align with ecological gradients (urbanisation, fragmentation, microhabitats) rather than temporal effects. Notably, 2020 (forest sampling) was warmer in the Paris region than 2021 (park sampling, 2020: 14.3 °C, 2021: 12.9 °C. Source: opendata.paris.fr), yet we sampled more thermophilic species in parks, which strengthens our inference. Finally, both habitats were sampled in the same season during rainless days, with sufficient effort to capture nearly all expected species (Figure S3). Taken together, these points support that the observed differences reflect genuine ecological responses to urbanisation and fragmentation rather than temporal sampling artefacts.

Another potential bias is that the spatial distribution of quadrats differed between forests and parks. In forests, quadrats were regularly spaced (every 9 m) along a 100 m transect, whereas in parks they were opportunistically placed in the wooded parts where leaf litter occurred. Indeed, it is not possible to lay 100 m long straight transects in parks because the wooded parts are not large enough. We also tried to scatter the sampling protocol in parks to limit the effects of possible heterogeneities therein. All quadrats were nevertheless separated by at least 9 m, as in forests, hence we do not think that this difference in their spatial distribution can impact the results.

A further limitation of our study is the lack of direct microhabitat characterisation. While we hypothesised that microhabitat variation likely drives the species richness observed in parks, we did not measure these variables directly. Although such measurements would provide a clearer link to our observations and represent a compelling direction for future research, quantifying microhabitats for ants remains a challenge. Our limited knowledge of the specific ecological niches for many species makes selecting relevant parameters difficult. For our research questions, the Winkler extraction seemed the best method to sample the litter ant community specifically. It is not destructive (a prerequisite to work in urban parks), fast, and does not require a second visit to recover a trap (unlike pitfall trapping), and it samples a small surface of the entire habitat. It is not a selective method, but this is advantageous for ants as it allows sampling subordinate species. Indeed, selective methods such as baiting tend to sample only the dominant species. Other taxa captured during the sampling (mites, spiders, coleoptera, etc.) have been kept for analysis of the food web and competitive contexts.

### Regional-scale effects of urbanisation and habitat fragmentation on species composition

At a regional scale, comparing rural versus urban environments reveals no differences in the total number of species but a clear difference in community composition, with beta diversity dominated by its turnover component. This suggests that dispersal is low between the two environments, creating different communities adapted to each, and highlights species sorting as a strong regional-scale mechanism in this metacommunity. Indeed, if dispersal between forest and urban sites was high, we would expect either community composition to be similar (low beta diversity) or, if colonisation success differed between the environments, that the communities of one environment would be a subset of the communities of the other environment (high nestedness, Baselga, 2010). Differences in community composition between forests and parks can also reflect differences in fragmentation and shifts in ecological niches. In parks, wooded areas represent only a portion of the park and are often non-contiguous and surrounded by open habitats, allowing workers from these patches to forage in the sampled litter. In contrast, even the most fragmented forest patches were large enough to allow sampling away from the forest edge. Climate represents another niche axis: parks are warmer than forests due to the urban heat island effect, favouring warm-adapted species, as confirmed by our results. This pattern has been observed in other taxonomic groups, such as wild bees (Geppert et al., 2023) or spiders (Meineke et al., 2017), placing our results in a general trend across taxa. These trait syndromes, i.e., traits that frequently occur together across different species, have also been observed in an urban context for traits related to the mobility or dietary specialisation (Hahs et al., 2023). Our study did not differentiate urbanisation and fragmentation at the global scale, as both are here confounded. It would be interesting to disentangle these two factors by studying urban landscapes with low fragmentation and forest landscapes with high fragmentation.

More species are present per urban park than per forest. Several explanations can be proposed. First, as mentioned above, species sorting dominates, which requires intermediate dispersal, sufficient to reach suitable patches and habitat heterogeneity generating distinct niches for species to sort into. In this context, parks may host more ant species than forests primarily if they harbour a higher diversity of habitats and microenvironments. Dispersal within the city may in fact not be strongly limiting, particularly given frequent human-mediated transport of soil, plants and materials, and given that the species found in parks have already been filtered by urban conditions and may include good dispersers. We however stress that dispersal is not too high, as very high levels would create a global homogenisation that is not consistent with the patterns reported here. This distinction matters for interpreting the underlying mechanisms: if parks are richer mainly due to habitat heterogeneity, species sorting would operate locally within parks in response to fine-grained environmental variation, rather than reflecting dispersal limitation at the regional scale. This is consistent with the trait differences we observe in urban communities (higher thermophily, tendency towards monogyny): monogynous species disperse via winged queens over potentially long distances, which argues against dispersal being strongly limiting for the species able to persist in the city, and suggests urban ant communities may instead be filtered towards good dispersers. This is not incompatible with previous findings that dispersal and competitive ability trade off within a single ant species under urbanisation (Finand et al., 2023). Dispersal ability may determine which species can colonise urban parks in the first place, even if it also declines within persisting populations. Disentangling these within- and between-species processes would require direct data on dispersal traits and habitat heterogeneity across parks.

Although our sampling within parks targeted shaded litter patches under bushes or trees to mimic forest conditions, they were often surrounded by more open and trampled areas, potentially creating a range of microhabitats. This aspect, not fully quantifiable in our study due to a lack of direct microhabitat characterisation, could contribute to higher species richness in parks. However, if parks were richer simply because of a higher microhabitat heterogeneity, we would expect to find a high nestedness. Instead, our data reveal a near-complete turnover in species composition between forests and parks, showing that parks do not merely add species but rather replace forest species with species better adapted to urban conditions. At a regional biogeographic scale, warmer climates are generally associated with higher species richness. By locally emulating warmer conditions through the urban heat island effect, parks may draw on a larger regional pool of thermophilic colonisers than surrounding forests, independently of any local filtering process determining which of these colonisers ultimately establish and persist. This is consistent with our trait analysis, which shows more warm-adapted species in parks. Another possibility is that species richness in parks might be “inflated” by continuous colonisation processes such as soil inputs during park maintenance or transport of material within urban areas. These hypotheses would implicitly question our view of forests being good patches compared to parks. An alternative explanation would be that forests are (or have been until relatively recently) well-connected and form a metacommunity of their own. If so, high dispersal among forests could homogenise ant forest communities through mass effects, leading to a depressed lower alpha diversity reflecting partial homogenisation (Leibold et al., 2004; Mouquet & Loreau, 2003). In contrast, when dispersal is low, as may be the case between parks, competition occurs independently within each park, resulting in different communities, hence higher alpha and gamma diversities. This increases turnover between patches as observed in our data. This is likely given that parks were made for human leisure and differ more from one another than natural forests do.

### Local-scale effects of habitat amount within parks and forests

Our observations of local community compositions support the idea of strong mass effects among forests. Indeed, there is no difference in species richness and species composition among forests, even for those with the lowest habitat amount. This suggests a high degree of dispersal between forest sites, which homogenises the communities. This is plausible because the matrix separating forests is less hostile than the one separating parks. Indeed, field hedges, roadsides and private gardens may connect forests by a network of trees and shrubs producing leaf litter and facilitating the dispersal of ants. An alternative, not mutually exclusive, explanation is also plausible. Forest patches are relatively large and environmentally homogeneous compared to parks. Under these conditions, even without strong dispersal, low extinction rates in large patches could allow species sorting to remain close to “optimal”, with similar, and likely temporally stable, community composition across patches despite variation in patch size. Both mechanisms would predict the homogeneity we observe, and distinguishing between them would require additional data on colonisation-extinction dynamics or dispersal rates between forest sites. Moreover, forests may be more similar to one another than parks. The former evolved from natural forests and have been managed largely for similar usages (e.g. wood production, leisure, hunting), whereas the latter are artificial and have to some extent been conceived to differ from one another to provide varied scenic environments. This could be confirmed by a more systematic study of biotic and abiotic environmental variations. If so, such homogeneity could limit the number of local niches, hence species sorting.

In urban areas, on a park scale, species richness is generally lower in low habitat amount parks, and community composition differs due to high species turnover. This pattern aligns with the classical species-area relationship, which links larger areas to higher diversity (Preston, 1962), potentially reflecting stronger competitive exclusion due to fewer available niches in smaller parks (Leibold et al., 2004; Mouquet & Loreau, 2003). In smaller parks, competitive exclusion may be more pronounced, and extinction rates are likely higher, as only the most competitive species persist, while low dispersal limits the arrival of new species. This limited dispersal is exacerbated by the hostile urban matrix surrounding these parks, which restricts movement and colonisation. Smaller parks likely receive fewer human-mediated dispersal events in absolute terms (e.g., less plant or soil material introduced overall). Consequently, these combined niche and dispersal effects contribute to lower alpha diversity and higher turnover between smaller parks. However, we found no correlation between species richness and park isolation (proximity index). This is apparently contradictory, but could be explained by the low variability in park isolation we have in the Parisian context. These patterns are consistent with other studies, including other ant communities. Smaller parks in highly urbanised zones tend to harbour fewer ant species than those in peri-urban, less isolated areas (Pacheco & Vasconcelos, 2007), while within cities, larger parks generally support greater alpha diversity (Carpintero & Reyes-López, 2014). This trend extends to other ecosystems and taxa. For example, forests show similar patterns (Leal et al., 2012). Other studies on isolated zooplankton communities in controlled environments also report reduced diversity due to limited dispersal (Steiner & Asgari, 2022), mirroring the low mass effects in our urban ant metacommunity.

Park management practices may affect urban biodiversity (Trigos-Peral et al., 2020). Note that this administrative management (e.g. mowing frequency, pruning, litter removal) is distinct from the landscape design heterogeneity discussed above. Even though most of our parks are managed by the same authority (the City of Paris), which may homogenise routine maintenance practices to some extent, this does not preclude the parks from differing markedly from one another in their original design, vegetation composition, and scenic intent. Management is further delegated at the city-district level, which could still introduce some variability in practices, potentially influencing our ant communities.

An interesting recent debate proposes to distinguish the effects of habitat amount vs. fragmentation *per se* (Fahrig, 2013, 2017; Riva et al., 2024), defined at a landscape scale based on the degree of subdivision (or breaking apart) of a given amount of habitat into several patches. We here chose to use habitat amount at the local scale, while our use of fragmentation is broader and encompasses habitat loss and isolation metrics. Indeed, when habitat loss and fragmentation are often concomitant through the urbanisation process, as is the case here, the distinction between the processes becomes largely semantic.

### Effects of urbanisation and habitat fragmentation on species traits

Beyond the turnover in species composition, we highlight that phenotypic traits are deterministically associated with community variations. Urban species are more thermophilic, and tend to be more often monogynous, than forest species. The high abundance of thermophilic ants in urban areas aligns with numerous studies indicating that cities, such as Paris, are subject to the urban heat island effect (Cantat, 2004; Lemonsu et al., 2015). Heat-susceptible species may disappear, and thermophilic species may dominate communities in cities. Parr and Bishop (2022) suggest that temperate ant species may be less vulnerable to climate warming than tropical species, and in some cases might even benefit from rising temperatures. However, our findings, showing a near-complete replacement of forest ant communities by species with more southern affinities in urban parks, do not fully support this idea. On the contrary, although temperature is not the only factor differing between urban and rural forests, our results suggest that our temperate ant communities living in forests may be negatively impacted by rising temperatures. The prevalence of thermophilic species in urban areas has been documented not only in ants but also in other groups, such as carabid beetles (Piano et al., 2017) and moths (Franzén et al., 2020). More broadly, the process of thermophilisation, where communities shift towards species that prefer warmer conditions due to rising temperatures, has also been observed in non-urban ecosystems, including fish (Daufresne et al., 2004) and plant communities (Feeley et al., 2020). This result is particularly relevant in the context of climate change, as temperatures are expected to rise in the coming decades, leading to significant ecological consequences (McCarty, 2001). Cities offer a valuable natural laboratory for studying the impacts of climate change, with urban communities serving as ideal experimental units to investigate these effects. While urban areas might serve as reservoirs for future biodiversity under changing climates, they will likely become increasingly inhospitable for many species as climate change progresses. An alternative, non-mutually-exclusive explanation is that the shift in thermophilic species reflects differences in habitat structure. Wooded patches within parks are smaller and more exposed to open, sun-lit surroundings than forest interiors, and species adapted to open or edge habitats are typically more thermophilic by nature than strict forest specialists. Distinguishing between these explanations would require direct microclimate measurements at each site. Both mechanisms (the urban heat island effect and the reduction of park wooded patches to small, edge-dominated fragments) are direct consequences of urbanisation and fragmentation and are therefore not fully separable in our design.

While urban thermophily was expected, given the heat island effect in cities, the tendency towards monogyny was not. Monogyny is somewhat linked to dispersal capacity (Keller, 1991; Sundström et al., 2005). Monogynous colonies are often founded by winged queens that disperse alone by flying over long distances. In contrast, polygynous colonies are either founded by related non-flying queens that disperse on foot over short distances with workers, or arise secondarily by non-flying queens that remain in their natal colony. Theoretically, habitat fragmentation can select for high dispersal (i.e. winged queens), for instance, when dispersal is traded off with competition capacity (Finand, Monnin, et al., 2024; Tilman, 1994), when fragmentation leads to high kin competition in remaining patches, or to avoid inbreeding (Cote et al., 2017; Gandon, 1999; Hamilton & May, 1977). Some empirical studies support these theoretical expectations, including in ants. For instance, at the intraspecific level, colonisation ability can be in a trade-off with competition in an ant species (Finand, Loeuille, et al., 2024). This selection for high dispersal in a fragmented context could explain, as a byproduct, that city parks harbour more monogynous species because polygynous species disperse less. However, studies focusing on dispersal *per se* have shown opposite results, with habitat fragmentation in cities selecting for lower dispersal (Cheptou et al., 2008; Finand et al., 2023). Indeed, while habitat fragmentation usually favours high dispersal, it may have the opposite effect under some circumstances (e.g. highly hostile matrix increasing dispersal costs, Bonte et al., 2012; Hastings, 1983). A second explanation is that monogynous species can be considered as pursuing spatial bet-hedging because flying queens can access and distribute the risks on more patches than walking queens. Indeed, a colony producing flying queens can attempt to establish new colonies on many patches. In contrast, a colony producing apterous queens produces a few only, as each requires the allocation of workers, and they have restricted dispersal, hence it has few establishment attempts. A bet-hedging strategy is selected under temporally variable environments (Slatkin, 1974). City patches may be more variable, due to many disturbances associated with human activities, and parks being surrounded by unfavourable environments, or due to extreme climatic conditions being less buffered. Having winged queens will increase the probability of having some offspring in a favourable environment. A third hypothesis is that many urban parks could be relatively new, particularly regarding ants that have a longer generation time than most other insects. The first species colonising a new patch are the most dispersive ones, which in ants have winged queens and thus are monogynous. Dispersal limitation also likely lasts in this highly fragmented context. Forests being much older, the less dispersive but more competitive species had the time to colonise them and progressively exclude better colonisers. All these hypotheses are not exclusive. Other taxa have shown this shift in community composition with more dispersive species in the city, like carabid beetles (Piano et al., 2017). Our data show a trend towards monogyny being more abundant in urban parks than forests, and future works may seek to test whether this is indeed the case. Population genetic analyses could also help determine whether dispersal is indeed higher for monogynous species in urban parks, and more broadly, quantify actual dispersal rates between sites.

## Supporting information

Sup_mat

## Acknowledgements

We would like to thank the City of Paris and the French National Museum of Natural History for allowing us to sample the parks of the city. We also thank Xavier Espadaler and Bernard Kaufmann for their help in the identification of some species. We thank recommender Paul Savary as well as reviewers Joan Casanelles-Abella, Nicolas Deguines and Audrey Muratet for their comments on earlier versions of the manuscript

## Conflict of interest

The authors declare no conflict of interest.

## Notes

https://doi.org/10.5281/zenodo.14792980

### Competing Interest Statement

The authors have declared no competing interest.

### Summary of Updates

New version of the manuscript after taking into account the comments of PCI ecology

