## Supplementary material for "Urbanisation and habitat loss favour thermophilic and monogynous ant species": Sup_mat

a.

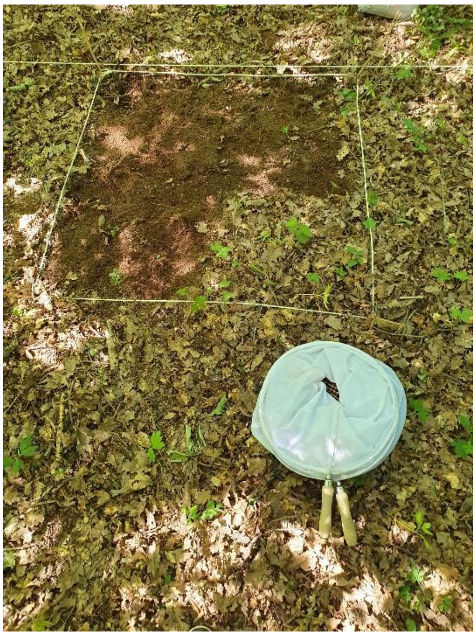

b.

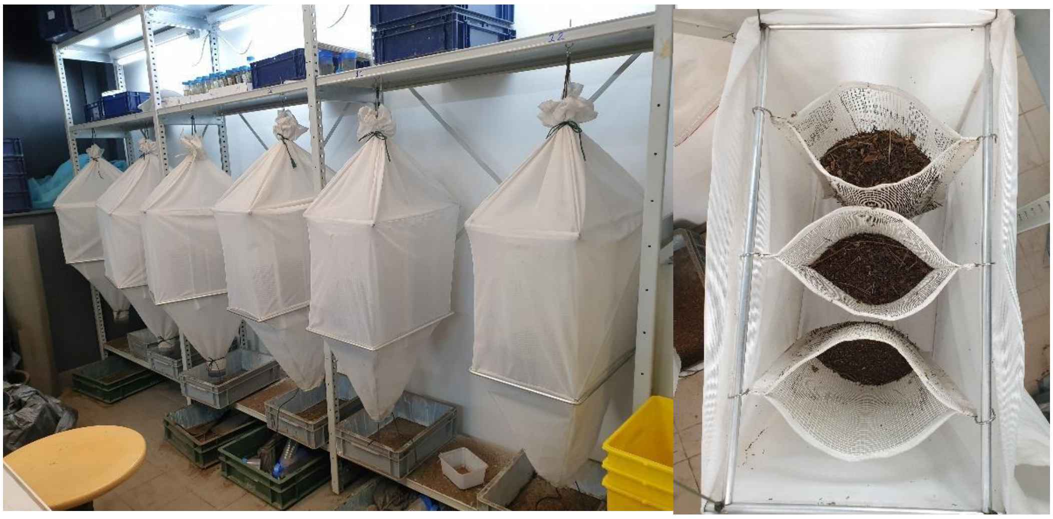

**Figure S1:** Pictures of (a) a quadrat being sifted and (b) the Winkler devices in the laboratory.

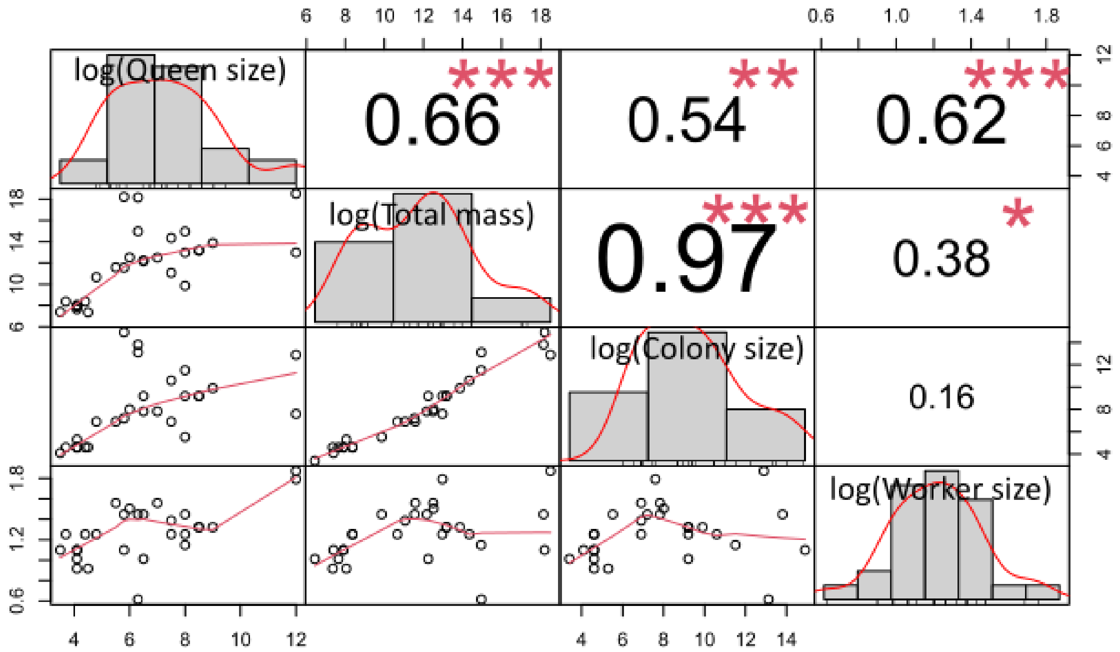


**Figure S2:** Correlation chart between the different size measurements at the species level. Diagonal histograms represent the distribution of each variable. The bottom of the diagonal represents the bivariate scatter plots and the top of the diagonal represents the value of the correlation. Statistical significance is calculated using a Pearson test represented by the red stars (* p-value < 0.05, ** p-value < 0.01, *** p-value < 0.001). The figure is done with the R package *PerformanceAnalytics.* We observe a strong correlation between all the size measurements.

**a.**

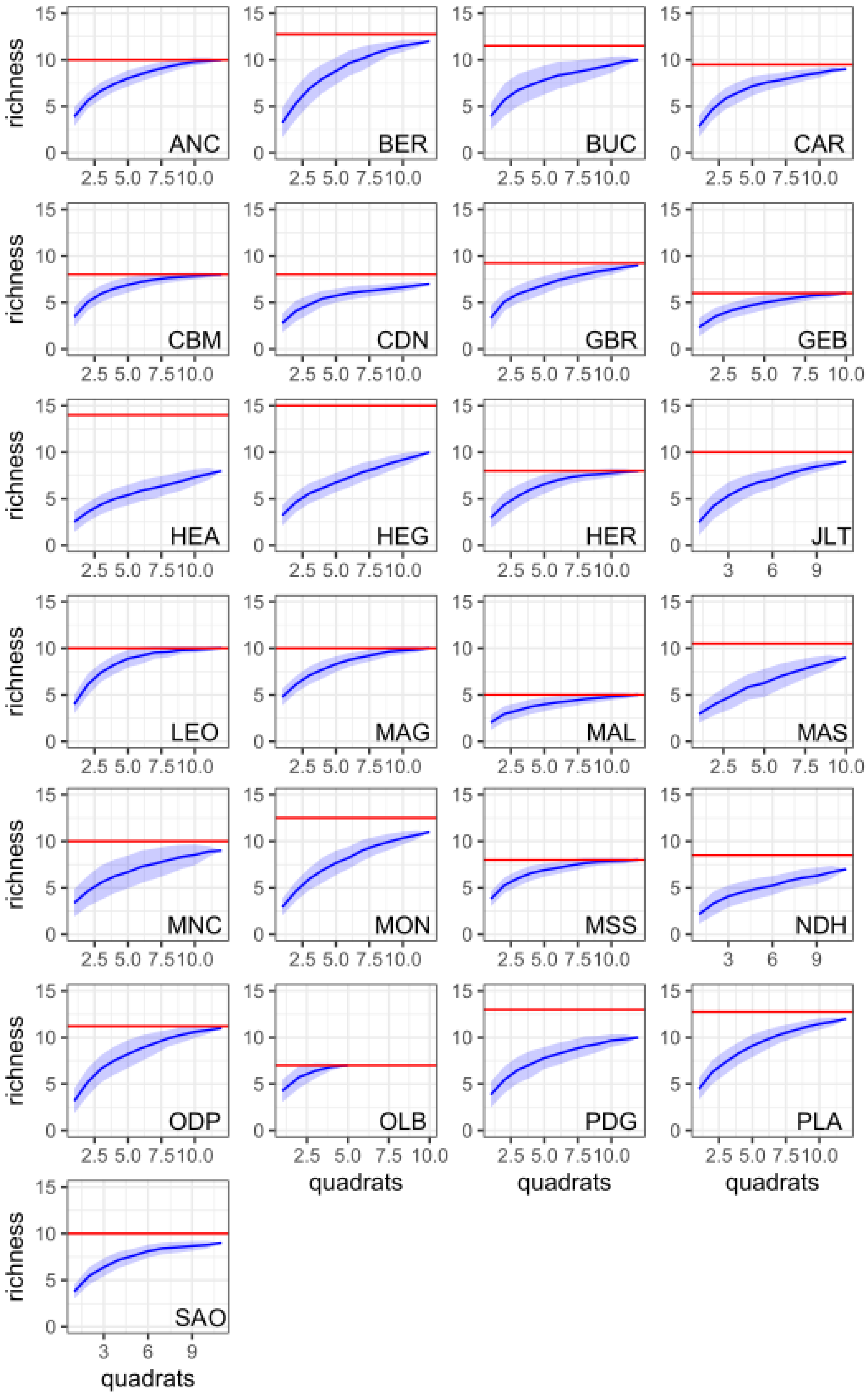
richness
quadrats
ANC
BER
BUC
CAR
CBM
CDN
GBR
GEB
HEA
HEG
HER
JLT
LEO
MAG
MAL
MAS
MNC
MON
MSS
NDH
ODP
OLB
PDG
PLA
SAO

b.

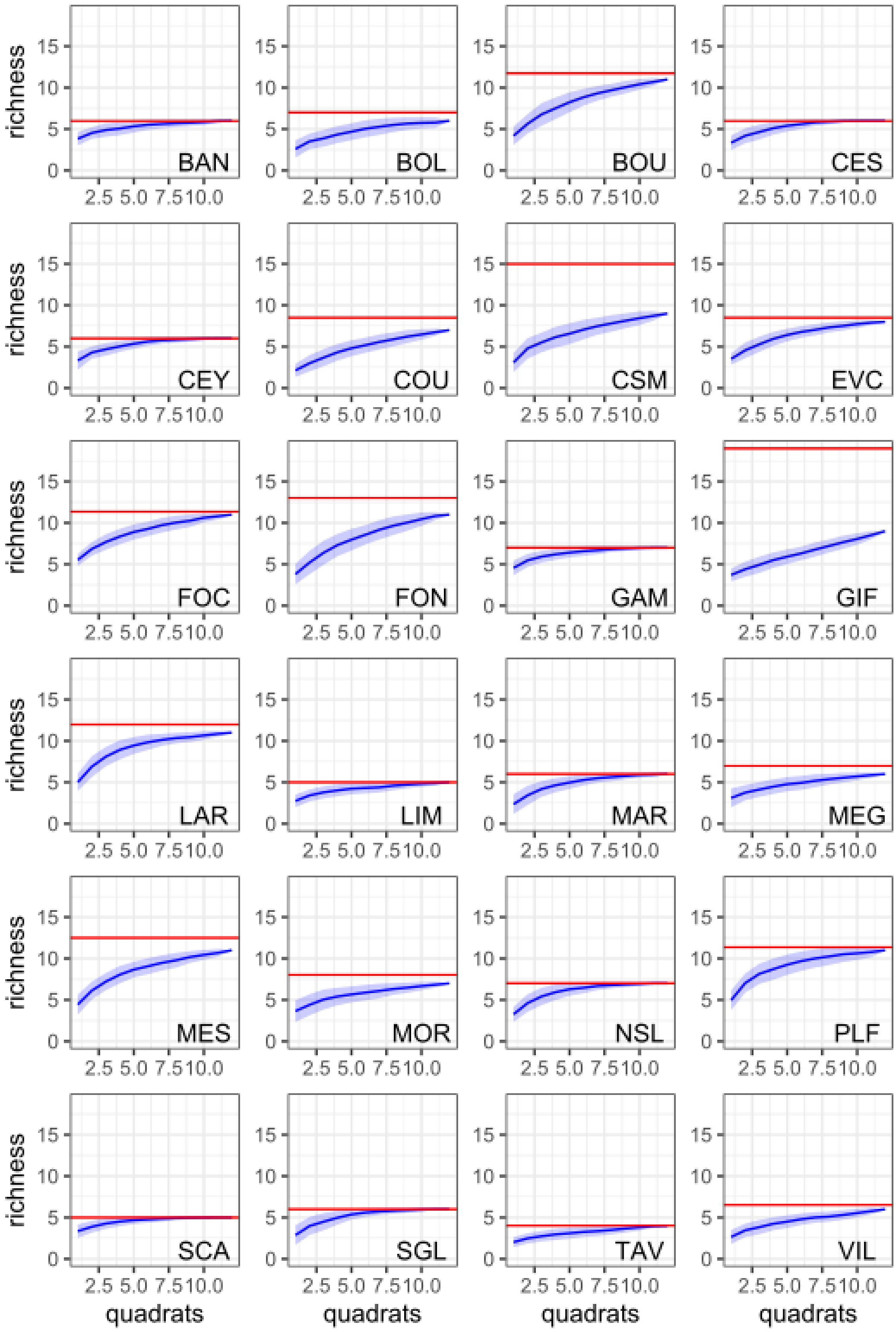


**Figure S3:** Rarefaction curve for urban sites (a) and forest sites (b). The letters inside the graph are the site references. The red line is the estimated number of species using the Chao method.

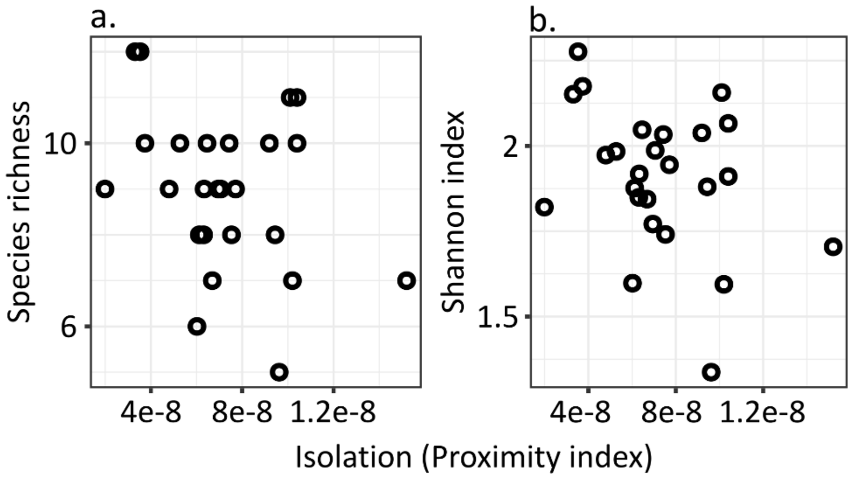


**Figure S4:** Relation between species richness (a) and Shannon index (b), and the isolation of the parks (proximity index, % of green area (other parks) weighted by its distance within 1 Km around the focal park boundaries). No significant differences are observed in species richness and Shannon index depending on the isolation (Species richness: t value = -1.63, p = 0.12; Shannon index: t value = -1.85, p = 0.08).

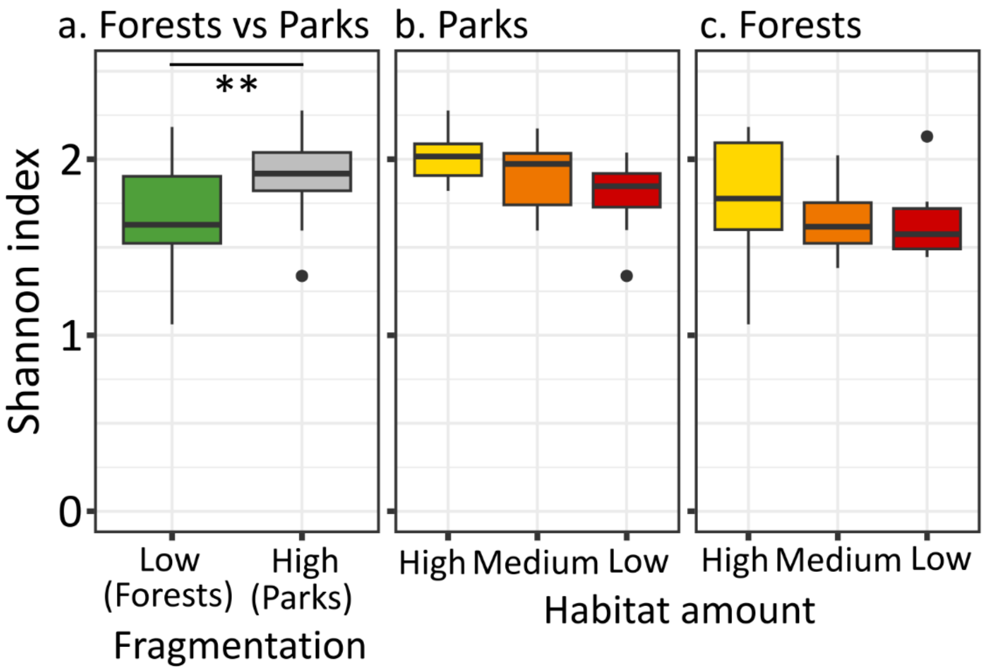


**Figure S5:** Shannon index between forests and parks (a), between parks of different habitat amount (b), and between forests of different habitat amount (c). Significant differences are observed in the Shannon index between forests and parks (F value = 7.89, p = 0.007, $R^2$ = 12.6%), a marginal effect within the parks (F value = 2.83, p = 0.081), and no effect within the forests (F value = 0.488, p = 0.62).

**Table S1:** Site characteristics. The parks have been classified into habitat amount categories depending on surface: high (80,000 to 250,000 m²), medium (5,000 to 12,000 m²), and low habitat amount (500-2,000 m²). Forests have been selected based on the habitat amount (percentage of forest within 1 Km radius): high (75 to 95% of forest cover), medium (30 to 70% of forest cover), and low habitat amount (less than 5% of forest cover). Isolation is based on the proximity index: the percentage of green areas weighted by their distances within a 1 Km buffer around park boundaries.

| Site | Habitat | Habitat amount | Isolation | Latitude | Longitude |
|---|---|---|---|---|---|
| ANC | PARK | HIGH | 6.46E-08 | 48.841253 | 2.274779 |
| BER | PARK | HIGH | 3.53E-08 | 48.835645 | 2.381719 |
| BUC | PARK | HIGH | 5.27E-08 | 48.879555 | 2.382098 |
| CAR | PARK | MEDIUM | 7.06E-08 | 48.891660 | 2.331571 |
| CBM | PARK | HIGH | 6.12E-08 | 48.891320 | 2.313937 |
| CDN | PARK | MEDIUM | 1.52E-07 | 48.853077 | 2.270599 |
| GBR | PARK | HIGH | 6.33E-08 | 48.831523 | 2.299992 |
| GEB | PARK | LOW | 6.02E-08 | 48.839894 | 2.297988 |
| HEA | PARK | MEDIUM | 7.53E-08 | 48.831372 | 2.370190 |
| HEG | PARK | LOW | 1.04E-07 | 48.851365 | 2.361816 |
| HER | PARK | LOW | 6.31E-08 | 48.827855 | 2.352126 |
| JLT | PARK | LOW | 7.71E-08 | 48.889220 | 2.338124 |
| LEO | PARK | MEDIUM | 3.74E-08 | 48.886116 | 2.353361 |
| MAG | PARK | MEDIUM | 7.43E-08 | 48.861687 | 2.379144 |
| MAL | PARK | LOW | 9.62E-08 | 48.882002 | 2.283976 |
| MAS | PARK | LOW | 6.95E-08 | 48.891951 | 2.361182 |
| MNC | PARK | HIGH | 1.99E-08 | 48.879372 | 2.308870 |
| MON | PARK | HIGH | 1.04E-07 | 48.821991 | 2.337792 |
| MSS | PARK | MEDIUM | 9.44E-08 | 48.870636 | 2.396616 |
| NDH | PARK | MEDIUM | 1.02E-07 | 48.852250 | 2.294367 |
| ODP | PARK | MEDIUM | 1.01E-07 | 48.835143 | 2.336580 |
| OLB | PARK | LOW | 6.69E-08 | 48.856201 | 2.380958 |
| PDG | PARK | LOW | 9.19E-08 | 48.850952 | 2.313205 |
| PLA | PARK | HIGH | 3.31E-08 | 48.843222 | 2.359509 |
| SAO | PARK | MEDIUM | 4.80E-08 | 48.887704 | 2.293356 |
| BAN | FOREST | LOW | - | 49.126160 | 1.802070 |
| BOL | FOREST | LOW | - | 48.874700 | 2.917940 |
| BOU | FOREST | MEDIUM | - | 48.422100 | 2.262050 |
| CES | FOREST | MEDIUM | - | 48.555610 | 2.589450 |
| CSM | FOREST | MEDIUM | - | 48.850050 | 2.590580 |
| CEY | FOREST | HIGH | - | 48.602920 | 1.902140 |
| COU | FOREST | LOW | - | 48.640940 | 2.958760 |
| EVC | FOREST | LOW | - | 48.621420 | 2.401270 |
| FON | FOREST | HIGH | - | 48.415790 | 2.623140 |
| FOC | FOREST | HIGH | - | 49.156570 | 2.608040 |
| GAM | FOREST | HIGH | - | 48.757850 | 1.714860 |
| GIF | FOREST | MEDIUM | - | 48.694020 | 2.137140 |

| | | | | | |
|---|---|---|---|---|---|
| LAR | FOREST | HIGH | - | 48.493840 | 2.653260 |
| LIM | FOREST | MEDIUM | - | 48.646550 | 2.087960 |
| MAR | FOREST | MEDIUM | - | 48.659130 | 2.214520 |
| MES | FOREST | LOW | - | 48.360130 | 2.235410 |
| MEG | FOREST | MEDIUM | - | 49.025280 | 2.766200 |
| MOR | FOREST | MEDIUM | - | 48.924180 | 1.934830 |
| NSL | FOREST | HIGH | - | 48.244550 | 2.790140 |
| PLF | FOREST | HIGH | - | 48.691940 | 1.797210 |
| SCA | FOREST | LOW | - | 49.064840 | 1.749630 |
| SGL | FOREST | HIGH | - | 48.966230 | 2.123320 |
| TAV | FOREST | HIGH | - | 49.036690 | 2.262790 |
| VIL | FOREST | HIGH | - | 48.803560 | 2.875580 |
